# Chromosome-scale genome assembly of the Hunt bumble bee, *Bombus huntii* Greene, 1860, a species of agricultural interest

**DOI:** 10.1101/2024.04.30.591905

**Authors:** Jonathan Berenguer Uhuad Koch, Sheina B. Sim, Brian Scheffler, Jeffrey D. Lozier, Scott M. Geib

## Abstract

The Hunt bumble bee, *Bombus huntii*, is a widely distributed pollinator in western North America. The species produces large colony sizes in captive rearing conditions, experiences low parasite and pathogen loads, and has been demonstrated to be an effective pollinator of tomatoes grown in controlled environment agriculture systems. These desirable traits have galvanized producer efforts to develop commercial *B. huntii* colonies for growers to deliver pollination services to crops. To better understand *B. huntii* biology and support population genetic studies and breeding decisions, we sequenced and assembled the *B. huntii* genome from a single haploid male. High-fidelity sequencing of the entire genome using PacBio, along with HiC sequencing, led to a comprehensive contig assembly of high continuity. This assembly was further organized into a chromosomal arrangement, successfully identifying 18 chromosomes spread across the 317.4 Mb assembly with a BUSCO score indicating >98% completeness. Synteny analysis demonstrates shared chromosome number (*n* = 18) with *B. terrestris*, a species belonging to a different subgenus, matching the expectation that presence of 18 haploid chromosomes is an ancestral trait at least between the subgenera *Pyrobombus* and *Bombus sensu stricto*. In conclusion, these assembly outcomes, alongside the minimal tissue sampled destructively, showcase techniques for producing efficient, comprehensive, and continuous genome arrangements.

## Introduction

Bumble bees (Hymenoptera: Apidae, *Bombus* Latreille, 1802) are significant pollinators of flowering plants, resulting in at least seven species being subjected to domestication by commercial enterprises for crop pollination since the mid 1980s (Velthuis and van Doorn 2006). Bumble bees are most heavily used to pollinate tomatoes grown in controlled environment agriculture (Strange 2015). The most widely used bumble bee pollinator is arguably *Bombus terrestris* (Linnaeus, 1758), as their commercial use has expanded beyond its native range of Europe and Asia. To date, *B. terrestris* has been used to pollinate crops throughout Mesoamerica, South America, Japan, New Zealand, and Australia (Velthuis and van Doorn 2006). However, other countries, including the United States of America (USA) and Canada, have specific policies in place limiting the movement of non-native bumble bees across borders. As of 2022, *B. impatiens, B. huntii*, and *B. vosnesenskii* have been made available by major bumble bee producers across specific regions in North America.

The Hunt bumble bee, *B. huntii*, is native to western North America, and spans the countries of Canada, Mexico, and the USA (Koch *et al*. 2018). Population genetic analyses identified two to five genetic populations that correspond to past climate change and geographic variation (Koch *et al*. 2018). Relative to other North American bumble bees, *B. huntii* populations in the USA are an excellent candidate for commercial colony production due to high abundance in the wild, captive rearing success, low disease prevalence, and pollination effectiveness of tomatoes grown in controlled environment agriculture (Strange 2015; Koch *et al*. 2015; Baur *et al*. 2019; Mullins *et al*. 2020; Strange *et al*. 2023). However, unlike *B. impatiens* and *B. vosnesenskii*, there are no genomic resources for *B. huntii* (Sadd *et al*. 2015; Heraghty *et al*. 2020). Given the demonstration of genetic variation across populations, high-quality genome resources will be important for linking genotypes to phenotypes and ecotypes that will not only be useful for understanding the evolution and ecology of this species, but also may be used to improve its utility as a domesticated pollinator. For example, pollination effectiveness, production of sexually reproductive females and males, immunity to pathogens and parasites, gyne overwintering survival, and captive rearing success are all key traits that can greatly influence market availability, profitability, and sustainability of bumble bees used in agriculture (Thorp 2003; Velthuis and Van Doorn 2006) and could potentially be enhanced by understanding their underlying genetic basis.

In this paper, we present a near-chromosome-level assembly for *B. huntii* (iyBomHunt1.1), marking one of the initial genomes assembled and annotated under the Beenome100 initiative (http://beenome100.org). The primary goal of this consortium of scientists is to produce high-quality reference genomes representing more than 100 native bee species distributed in the USA. Employing PacBio HiFi generated data combined with a HiC library, we present an annotated genome assembly of *B. huntii* that is structured into 18 scaffolds mirroring 18 bumble bee chromosomes. The genome’s quality stands out favorably compared to previously sequenced bumble bee genomes and promises to be an asset for genomic investigations concerning this bee species, which holds ecological and agricultural significance.

## Methods & Materials

### Organism/strain origin and derivation

A male *B. huntii* specimen was used to develop the reference genome assembly. The specimen was collected by hand from a colony reared in captivity following bombiculture techniques described in Rowe et al. (2023). The foundress queen of the colony (mother of the male specimen) was collected in North Logan in Cache County, Utah (Coordinates: 41°45’54“N 111°48’48“W, 1400 m). The male specimen was flash frozen in liquid nitrogen and maintained at -80°C until they were shipped to the United States Department of Agriculture - Agricultural Research Service (USDA-ARS) – Pacific Basin Agricultural Research Center (PBARC) in Hilo, Hawaii.

### Sequencing methods and preparation details

Specimens were sent to the USDA-ARS PBARC to undergo DNA extraction and PacBio and HiC library preparation. Genomic DNA was extracted from a slice of abdominal tissue from a single *B. huntii* male (ToLID iyBomHunt1). The fresh or frozen tissue protocol of the Qiagen MagAttract HMW DNA Kit (Qiagen, Hilden Germany) was followed to obtain DNA that was sufficiently of high-molecular weight for PacBio sequencing. Isolated genomic DNA was purified using 2:1 polyethylene glycol with solid-phase reversible immobilization beads (DeAngelis *et al*. 1995). The resulting DNA was quantified using a dsDNA Broad Range (BR) Qubit assay (Thermo Fisher Scientific, Waltham, Massachusetts, USA) and the fluorometry function of a DS-11 Spectrophotometer and Fluorometer (DeNovix Inc, Wilmington, Delaware, USA). Purity was determined using OD 260/230 and 260/280 ratios from the UV-Vis spectrometer feature of the DS-11. The high-molecular weight DNA sample was then sheared to a mean size of 20 kb with a Diagenode Megaruptor 2 (Denville, New Jersey, USA) and the subsequent size distribution was assessed with an Agilent Fragment Analyzer (Agilent Technologies, Santa Clara, California, USA) using a High Sensitivity (HS) Large fragment kit. The PacBio SMRTBell library was prepared using the SMRTBell Express Template Prep Kit 2.0 (Pacific Biosciences, Menlo Park, California, USA). The prepared library was bound and sequenced on a Pacific Biosciences 8M SMRT Cell on a Sequel IIe system (Pacific Biosciences, Menlo Park, California, USA) at the USDA-ARS Genomics and Bioinformatics Research Unit in Stoneville, Mississippi. The run was performed with a 2-hour pre-extension followed by a 30-hour movie collection time. After sequencing, consensus sequences from the PacBio Sequel IIe subreads were obtained using the SMRTLink v8.0 software.

A HiC library was also prepared from a slice of abdominal tissue from the same male *B. huntii* used for HiFi sequencing using the Arima HiC kit (Arima Genomics, San Diego, California, USA) from crosslinked tissue prepared following the Arima HiC low input protocol. Following proximity ligation, DNA was sheared using a Bioruptor (Diagenode, Dennville, New Jersey, USA) and DNA fragments in the range of 200-600 bp were selected as the input for Illumina library preparation using the Swift Accel NGS 2S Plus kit (Integrated DNA Technologies, Coralville, Iowa, USA). Illumina sequencing (150 bp paired-end) was performed on a NovaSeq 6000 at the Hudson Alpha Genome Sequencing Center (Huntsville, Alabama, USA), and adapter trimming after sequence collection was performed using BaseSpace software (Illumina, San Diego, California, USA).

### Data analysis methods

Genome assembly methods largely follow Koch et al. (2023) but are briefly described below. First, HiFi reads containing artifact adapter sequences were removed using the program HiFiAdapterFilt v2.0 (Sim *et al*. 2022). The filtered HiFi reads were assembled into a contig assembly using HiFiASM v0.15.1-r329 (Cheng *et al*. 2021) using the default parameters and the output was converted to .fasta format using any2fasta (Seeman, 2018, https://github.com/tseemann/any2fasta). Scaffolding with HiC data was performed following the Arima Genomics mapping pipeline (Ghurye et al. 2019, https://github.com/ArimaGenomics/mapping_pipeline) and using the YaHS scaffolding software (Zhou *et al*. 2022) (https://github.com/c-zhou/yahs). The Arima Genomics mapping pipeline uses BWA mem (Li 2013) to align the paired Illumina reads separately to the HiFiASM contig assembly and uses the mapping pipeline script ‘filter_five_end.pl’ to retain reads mapped in the 5’ orientation. The individual read alignments were then processed with the ‘two_read_bam_combiner.pl’ script to produce a sorted and quality-filtered paired-end bam file. Picard Tools ‘MarkDuplicates’ (Picard Tools, 2019, https://broadinstitute.github.io/picard/) was used to remove PCR duplicates. The resulting .bam file and the HiFiASM contig assembly were input into YaHS for scaffolding using the ‘no contig error correcting’ option and converted to Juicebox (Durand *et al*. 2016) compatible files using the ‘juicer_pre’ function. Manual curation was then performed in Juicebox (v2.15) and edits were applied to the scaffold assembly using ‘juicebox_assembly_converter.py’ from Phase Genomics (https://github.com/phasegenomics/juicebox_scripts).

Local alignments to the nucleotide and protein databases were then used to assign the *B. huntii* contigs to a taxon using the rule ‘bestsumorder’ of blobtoolkit v.2.6.1 (Challis *et al*. 2020) which assigns contigs to a taxon first based on alignments to the NCBI nucleotide database and then followed by alignments to the protein database if there were no hits to the nucleotide database. Taxonomic assignment of assembled scaffolds was tertiarily conducted using NCBI Foreign Contamination Screen (FCS, https://github.com/ncbi/fcs/wiki) tool suite using the fcs-gx function which uses the genome cross-species aligner (GX) to identify contaminants of which there were none in the final assembly. Coverage per scaffold and contig record was calculated using minimap2 v2.2-r1101 (Li 2021).

The HiC scaffold assembly was assessed for completeness using a Benchmark of Universal Single Copy Orthologs (BUSCOs), with all relevant taxonomic databases for the genome (Eukaryota, Metazoa, Arthropoda, Insecta, and Endopterygota) and only the most derived database, Endopterygota for the protein set. *Ab initio* annotations on the scaffold assembly were performed using Metaeuk v.4.a0f584d (Levy Karin *et al*. 2020) for the Eukaryota, Arthropoda, Insecta, and Endopterygota odb10 databases and Augustus v3.4.0 (Stanke *et al*. 2008) were used to detect the Metazoa odb10 orthologs. Designation of genes as complete single copy, duplicated, fragmented, or missing were determined using BUSCO v5.2.2 (Manni *et al*. 2021) in ‘genome’ mode for the genome assembly and ‘protein’ for the annotated protein set. Identification for off-target (non-*B. huntii*) contigs in the assembly was performed by aligning all contigs to the NCBI nucleotide (NT) database (accessed 2022-02-14) using the ‘blastn’ function of BLAST+ v2.5.9+ (Camacho *et al*. 2009). The contigs were secondarily aligned to the UniProt protein database (accessed 2020-03) using Diamond (Buchfink *et al*. 2021).

Coverage, taxonomic assignment, and BUSCO results were aggregated using blobtoolkit and subsequently summarized using blobblurb v2.0.1(Sim 2022). Expected genome size was estimated using GenomeScope v2.0 (Ranallo-Benavidez *et al*. 2020) which uses k-mer frequency analysis of k-mer counts performed by KMC v3.2.1 (Kokot *et al*. 2017). Level of duplicate artifacts in the assembly was assessed using BUSCO results for both the genome and the protein set and using k-mer abundance in the raw HiFi reads relative to their representation in the final assembly as determined by K-mer Analysis Toolkit v2.4.2 (KAT) (Mapleson *et al*. 2017).

The *B. huntii* genome was submitted to the National Center for Biotechnology Information (NCBI) RefSeq (Rajput *et al*. 2019) for annotation using the NCBI Eukaryotic Genomic Annotation Pipeline v10.0. This method was selected to provide consistency and standardization in annotation methods with other bumble bee species (Sadd *et al*. 2015; Heraghty *et al*. 2020; Koch *et al*. 2023) and with the ongoing USDA-ARS sequencing initiatives AgPest100 and Beenome100. The annotation pipeline utilized *>*6 billion RNA sequencing reads from other closely related North American species in the subgenus *Pyrobombus* available on GenBank.

We assigned the annotated genes from *B. huntii* to orthologous groups along with publicly available gene annotations from *Apis mellifera* (NCBI GenBank accession: GCF_003254395.2), *B. (Bombus) terrestris* (NCBI RefSeq accession: GCF_910591885.1 and ToLID: iyBomTerr1), and *B. (Pyrobombus) hypnorum* Linnaeus, 1758 (NCBI GenBank and Ensembl accession GCA_911387925.2 and ToLID: iyBomHypn1 (Crowley and Sivell 2023)] using OrthoFinder (Emms and Kelly, 2019) with default parameters. We visualized the results from OrthoFinder using a riparian plot produced by GENESPACE (Lovell et al. 2022) to depict chromosomal distribution patterns and identify potential regions of the genome that are associated with genomic gaps, inversions, and translocations. Finally, we depicted shared sets of genes between taxa in an upset plot using UpSetR (Conway *et al*. 2017; Lex et al. 2014).

## Results and Discussion

### Genome Assembly

PacBio Sequel II HiFi and HiC sequencing of *B. huntii* produced sufficient data for highly contiguous contig assembly with chromosomal resolution. In the preliminary contig assembly, 55.2% (sum of reads = 183M) of the contigs did not match any taxonomic group (Figure 2A), potentially signaling a significant number of contaminants, which is not unexpected for abdominal tissue. The other contigs had BLAST hits to Arthropoda (37.2%), Streptophyta (7%), Nematoda (1%), and Ciliophora (<1%) (Figure 2A-B). Additional rounds of filtering with blobtools and alignment to HiC sequences, including the removal of “No Hit” sequences resulted in the removal of most non-Arthropod contaminants in the remaining scaffolds (Figure 2C-D). The 29 of the remaining 31 contigs taxonomically assigned as “non-Arthropod” collectively have 559 annotated genes and indicates spurious assignment as non-Arthropod.

**Figure 1.**
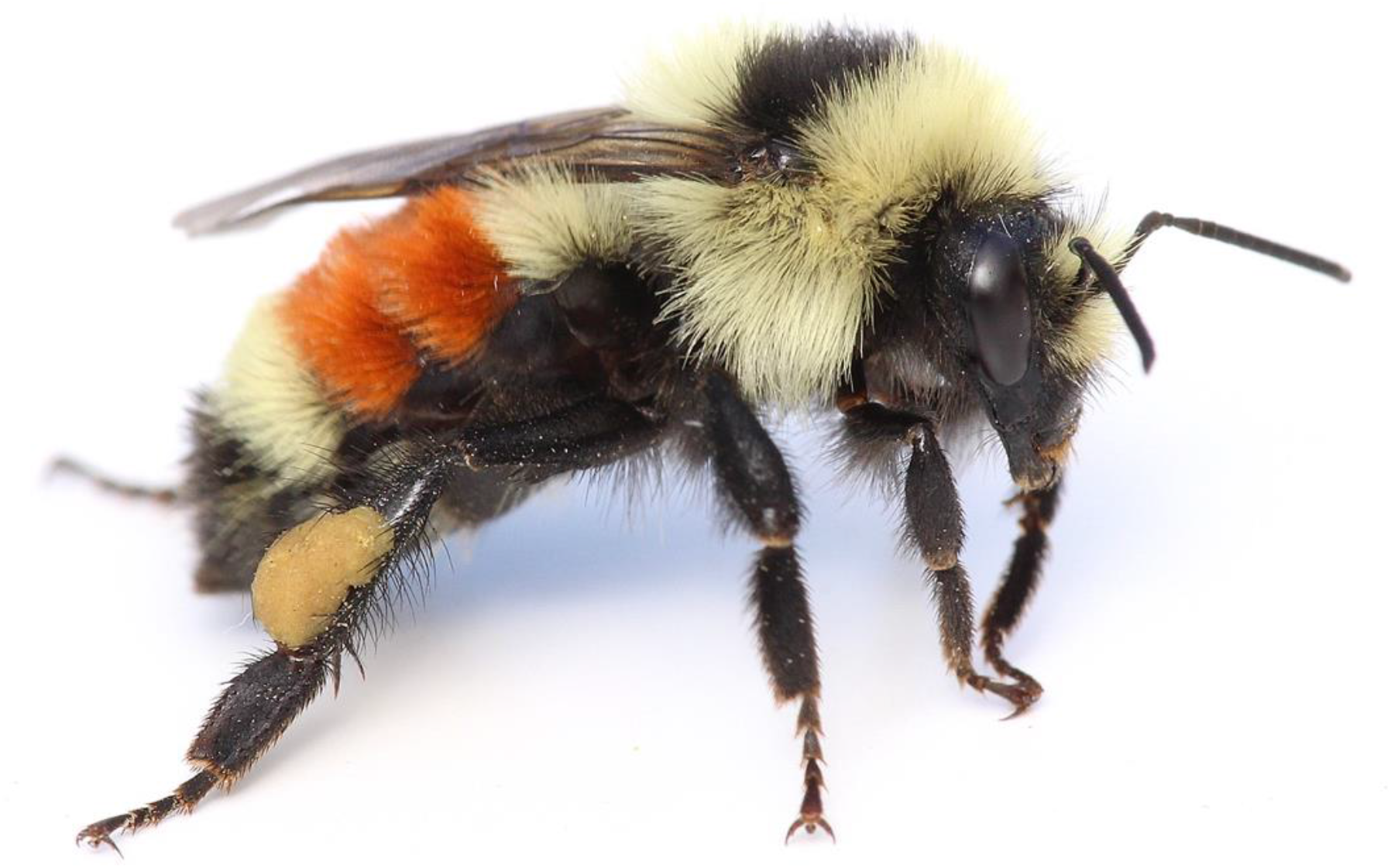
Lateral view of *Bombus huntii* gyne. Image by Joseph S. Wilson, with permission.

**Figure 2.**
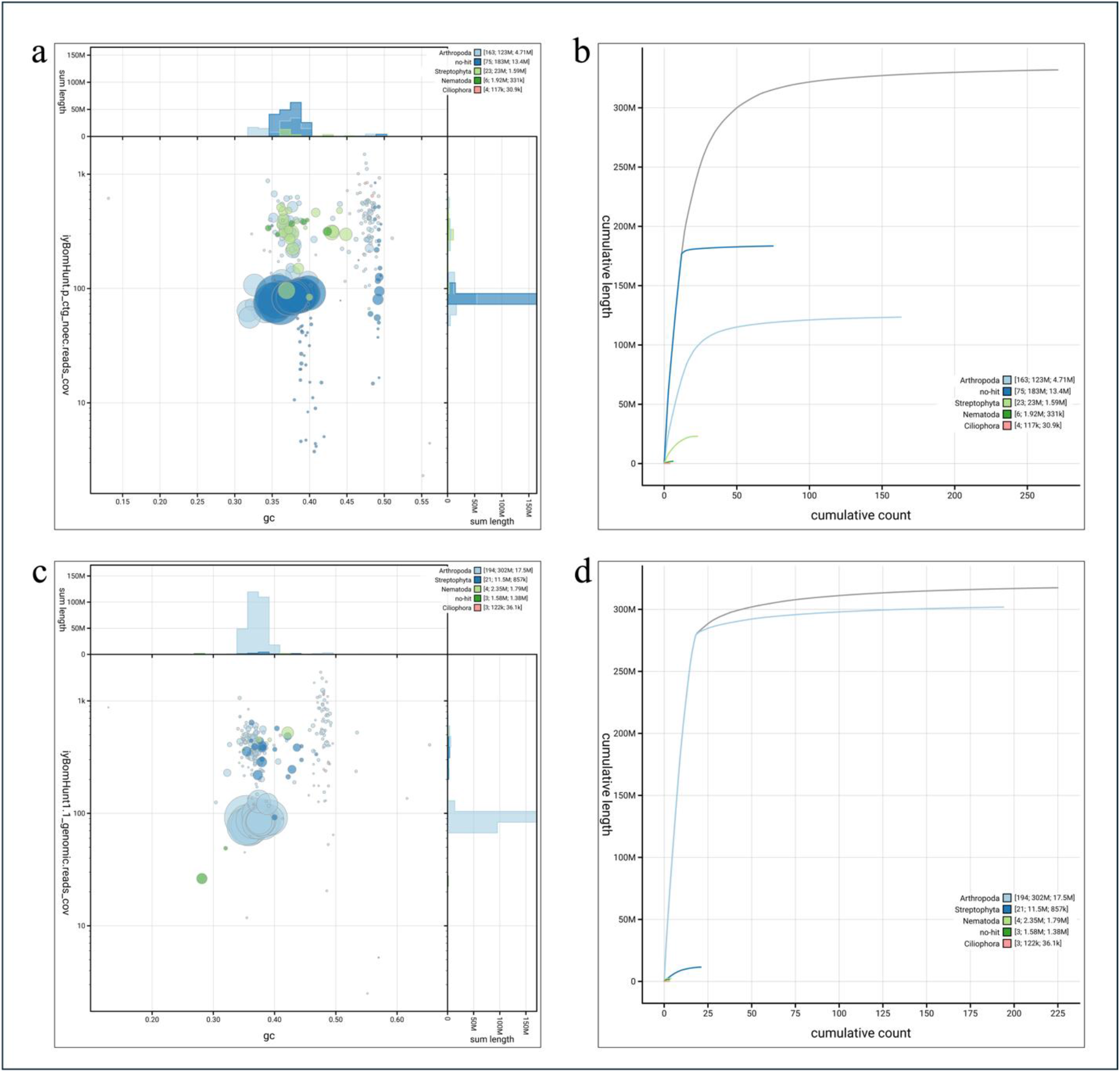
Blob plots from PacBio data showing read depth of coverage, GC content, and size of a) contigs, b) cumulative count and length of contigs per category, c) scaffolds, d) cumulative count and length of scaffolds per category.

The initial *B. huntii* assembly resulted in 225 scaffolds with a scaffold N50 of 17.5 Mb (Table 1) (Figure 3A). The assembly exceeds the minimum reference standard of 6.C.Q40 (>1.0 Mb contig and 10.0 Mb scaffold N50) identified by the Earth BioGenome Project (Lewin *et al*. 2018; Lawniczak *et al*. 2022). HiC contact mapping was able to recover 18 chromosome-length scaffolds (Figure 3B). The size of the scaffolded *B. huntii* assembly is 317.4 Mb (L50 = 8). Total haploid assembly size is estimated to be 288.73 Mb based on k-mer analysis with GenomeScope v2 (k-mer = 21, k-cov = 83) (Figure 3C). The assembly is hypothesized to have minimal error (=0.223%), likely due to low sequencing error and low repetitive content of sequences (=1.04%). The assembly is larger and more intact than the most recent *B. impatiens* Cresson, 1863 genome assembly (version BIMP_2.2) and other published genomes of closely related North American species in the subg. *Pyrobombus* [*B. bifarius* Cresson, 1878 = 266.8 Mb, *B. vancouverensis* Cresson, 1878 = 282.1 Mb, and *B. vosnesenskii* Radoszkowski, 1862 = 275.6 Mb] (Figure 3B), demonstrating the quality of the assembly compared to other *Bombus* genomes assembled without the advantages of PacBio and HiC data (Table 1). GC content of the *B. huntii* assembly is comparable to the species assemblies included in our study at 37% (average = 37.8% +/- 0.20%) (Figure 3B) (Table 1).

**Table 1.** Assembly statistics and BUSCO analysis for *B. huntii* in comparison to other Bombus genomes.

| Species | Assembly version | Len.<br>(Mb) | N50<br>(Mb) | L50 | No.<br>scaffolds | GC % | Complete<br>[single, duplicated] | Frag. | Miss. |
| --- | --- | --- | --- | --- | --- | --- | --- | --- | --- |
| *B. huntii* | iyBomHunt1.1 | 317.4 | 17.5 | 8 | 225 | 37 | 98.7% [98.4%,0.3%] | 0.20% | 1.20% |
| *B. bifarius* | Bbif_JDL3187 | 266.8 | 2.2 | 30 | 1,249 | 37.96 | 98.1% [97.7%,0.4%] | 0.60% | 1.30% |
| *B. vancouverensis* | Bvanc_JDL1245 | 282.1 | 3.06 | 24 | 1,162 | 38.02 | 98.4% [97.9%,0.5%] | 0.60% | 1.00% |
| *B. vosnesenskii* | Bvos_JDL3184-5_v1.1 | 275.6 | 2.83 | 24 | 1,429 | 37.93 | 98.2% [98.0%, 0.2%] | 0.60% | 1.20% |
| *B. impatiens* | BIMP_2.2 | 242 | 1.41 | 54 | 5,460 | 38 | 99.0% [98.8%, 0.2%] | 0.30% | 0.70% |
| *B. hypnorum* | iyBomHypn1.2 | 297.3 | 24.3 | 6 | 52 | 37.5 | 97.6% [97.4%, 0.2%] | 0.5% | 1.9% |
| *B. terrestris* | iyBomTerr1.2 | 393 | 14.6 | 10 | 249 | 38.5 | 98.9% [98.5%, 0.4%] | 0.10% | 1.00% |

**Figure 3.**
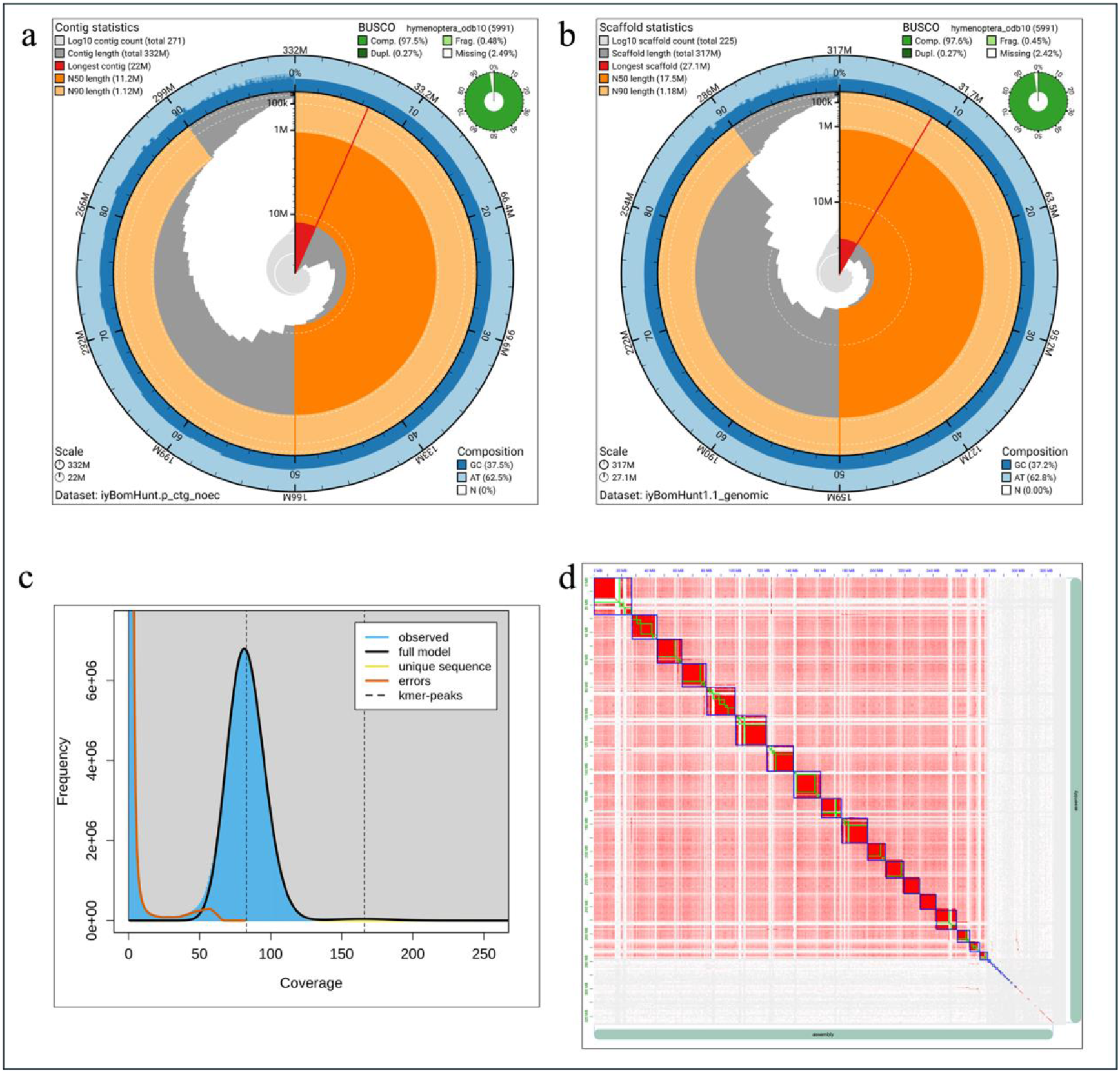
Snail plot visualization and genome assembly statistics for *B. huntii* with a) contig and b) scaffold resolution, c) Haploid assembly size estimated with GenomeScope v., and d) HiC contact map of *B. huntii* genome assembly. The HiC contact map demonstrates chromosomes ordered by size from left to right and top to bottom. The snail plot (contig and scaffold) is divided in one thousand size-ordered bins around the circumference. Each bin represents 0.1% of the assembly (contig = 332 Mb; scaffold = 317 Mb). The distribution of the scaffold lengths are shown in grey. The plot radius is scaled to the longest scaffold present in the assembly (contig = 22 Mb; scaffold = 27.1 Mb, both shown in red). The orange and pale-orange arcs show the N50 and N90 lengths (contig N50 = 11.2 Mb, contig N90 = 1.12 Mb; scaffold N50 = 17.5 Mb, scaffold N90 = 1.18 Mb). The blue and pale-blue ring around the outside of the plot is the distribution of GC, AT, and N percentages in the same bins as the inner plot. A summary of the BUSCO genes in the hymenoptera_odb10 dataset is shown in the top right of both contig and scaffold assemblies.

BUSCO scores also indicated a highly complete genome assembly (Figure 3B) (Table 1). The *B. huntii* assembly performed exceptionally better than all other published assemblies in the subg. *Pyrobombus*, with 98.7% of the 5,991-benchmarking universal single-copy orthologs represented in OrthoDB v1.10 Hymenoptera lineage dataset (hymenoptera_odb10). Most of the genes were single copy (98.4%) with 0.3% duplicated or 1.2% missing (Figure 3B) (Table 1).

### Genome Annotation and Synteny

In total, the NCBI Eukaryotic Genomic Annotation Pipeline predicted 15,072 genes and pseudogenes, of which 14,643 genes gave rise to 34,820 transcripts. The number of predicted genes in the *B. huntii* genome is greater than the number of genes predicted in *B. impatiens* and the *B. huntii* sister species *B. vosnesenskii* by 14.5% to 11.43%, although the number of predicted protein coding genes (11,088) is similar to other species (Table 2). Much of the difference in gene numbers could be attributed to the annotation of 3,554 non-coding genes in *B. huntii*, whereas an average of 1,884 non-coding genes were detected in the genome assembles of *B. bifarius, B. vancouverensis, B. vosnesenskii*, and *B. impatiens*. The elevated number of non-coding genes detected in the assembly may be due to the quality of the PacBio HiFi sequencing technology. Increased detection of non-coding genes in a bumble bee genome was also observed in the *B. affinis* genome assembly which used the same sequencing platform and bioinformatics methods described in this study (Koch *et al*. 2023). The verification, importance, and function of these non-coding genes in *Bombus* would benefit from further research.

**Table 2.**
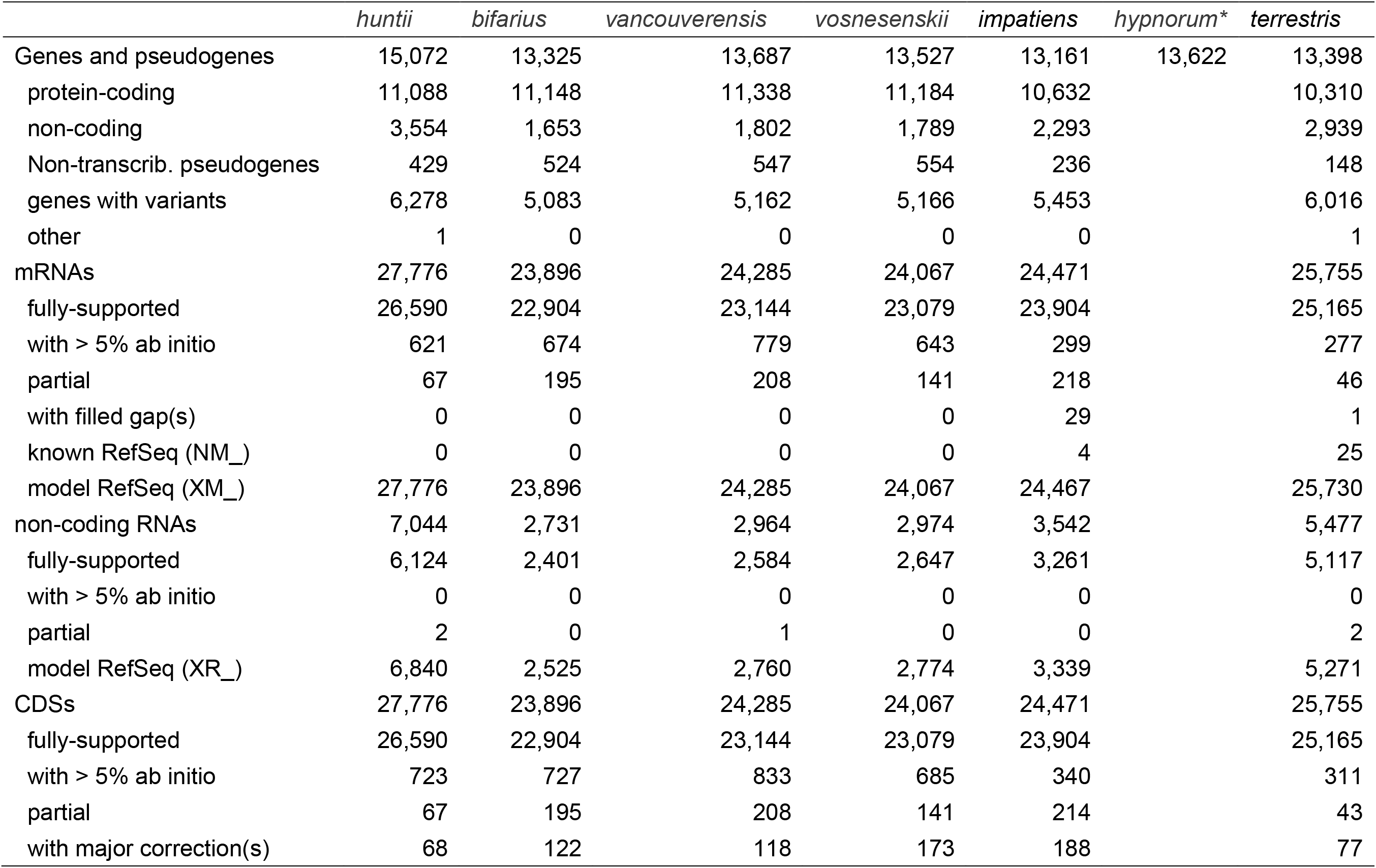
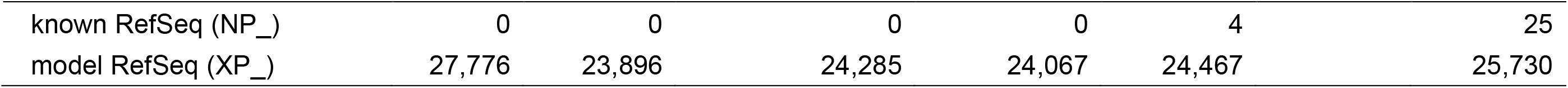
Annotation statistics from the NCBI eukaryotic genome annotation pipeline for *B. huntii* and other *Bombus* genomes available on NCBI. The B. hypnorum assembly was annotated with Ensemble rapid annotation pipeline (Crowley *et al*. 2023). Thus, the NCBI annotation statistics are not available for a comparative assessment with other focal *Bombus*.

We analyzed the synteny of *B. huntii* (*B. Pyrobombus*) with *B. terrestris (B. Bombus*), and *B. hypnorum* (*B. Pyrobombus*) as these species are represented by chromosomal resolution assemblies on NCBI. We also included *A. mellifera* in the synteny analysis as an outgroup for comparative purposes. The subgenus *Pyrobombus* (*i*.*e*., *B. huntii* and *B. hypnorum*) diverged from the subgenera *Bombus* (*i*.*e*., *B. terrestris*) and *Alpinobombus* around 19 mya during the Miocene (Hines 2008). Furthermore, the clade that includes *B. hypnorum* diverged from the clade the includes *B. huntii* around 15 mya, also during the Miocene (Hines 2008). Synteny analysis demonstrates that *B. huntii* and *B. terrestris* share the same number of linkage groups, hereafter identified as “chromosomes” (*n* = 18) (Figure 4). However, *B. hypnorum*, while in the same subgenus as *B. huntii*, has a reduced number of chromosomes (*n* = 12) (Figure 4). A chromosome number of 12 has also been reported in *B. perplexus* Cresson, 1863 through karyotyping (Owen *et al*. 1995). *B. perplexus* belongs to the same clade as *B. hypnorum* and is a sister taxon to *B. hypnorum* (Cameron *et al*. 2007; Hines 2008). Consistent with the hypothesis for *Bombus* as a whole (Owen *et al*. 1995), a haploid number of 18 chromosomes appears to represent the ancestral state shared between at least the subgenera *Bombus* and *Pyrobombus*, with 12 chromosomes to be a derived trait found in clade that includes *B. hypnorum* and *B. perplexus*. However, further sampling of taxa across *Pyrobombus* and *Bombus* will be required to confirm this hypothesis and investigate the potential timing and importance of this reduction in chromosome number in some *Pyrobombus*.

**Figure 4.**
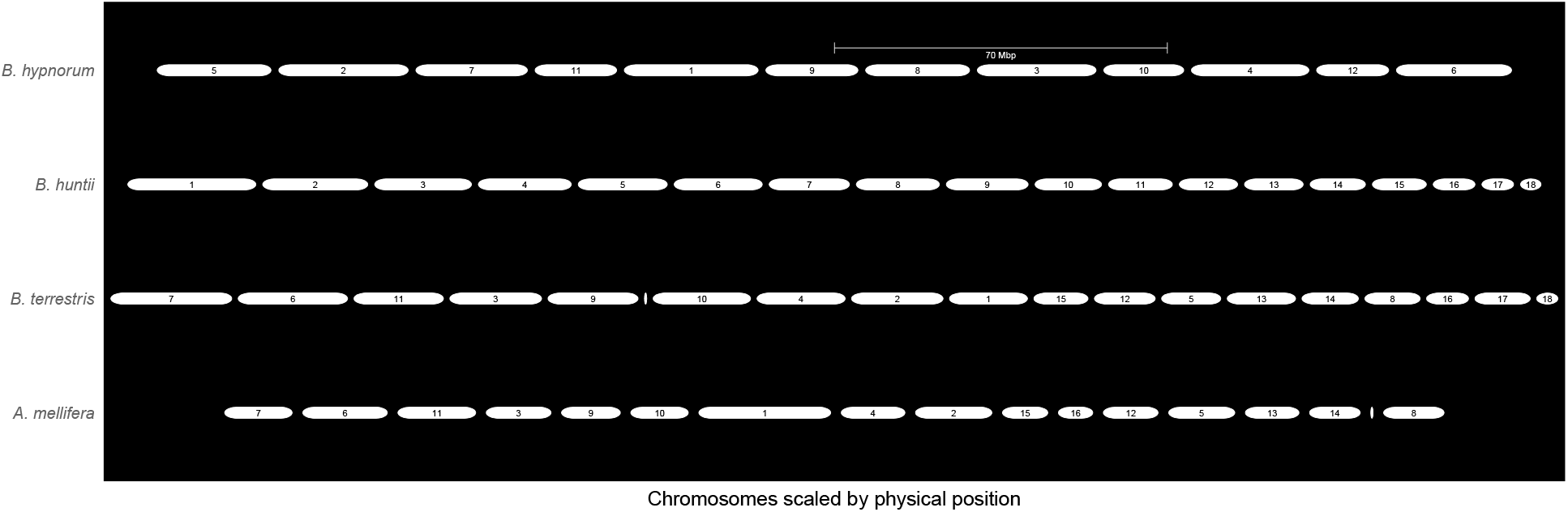
A GENESPACE-generated synteny map (bottom to top) of *A. mellifera* (outgroup), *B. terrestris, B. huntii* (study taxon), and *B. hypnoroum*. The white horizonal segments represent chromosomes. The colors (red to purple) represent the orthologous *B. huntii* chromosomes (1-18) and braids represent the syntenic blocks between bee genomes. X-axis positions are scaled by physical position.

Visualization of chromosome arrangements across *B. huntii, B. terrestris*, and *B. hypnorum* demonstrate several rearrangements including inversions, gaps, and repeats (Figure 4). Of course, the structural rearrangements could indicate assembly artifacts or genuine changes that occurred within the past 20 million years since the divergence from the common ancestors of *B. terrestris* with *B. huntii* and *B. hypnorum*, as well as between *B. huntii* and *B. hypnorum* (~15 mya). Interestingly, *B. hypnorum* does not appear to have lost major chromosomal regions with the reduction in chromosome number, with most *B. huntii* regions represented, indicating chromosome fusion events as the mechanism of this reduction (Owen *et al*. 1995). Finally, we compared the number of orthogroups shared across *B. huntii, B. hypnorum, B. terrestris*, and *A. mellifera*. We determined that 86% (*n* = 9,810) of the orthogroups identified through NCBI annotation were shared across the four focal species with 7,933 orthogroups found in single-copy and the remaining 509 orthogroups found in multiple copy in one or more of the four species. Furthermore, we identified 32 genes unique to *B. huntii* and 101 orthogroups shared between *B. huntii* and *B. hypnorum* (Figure 5).

**Figure 5.**
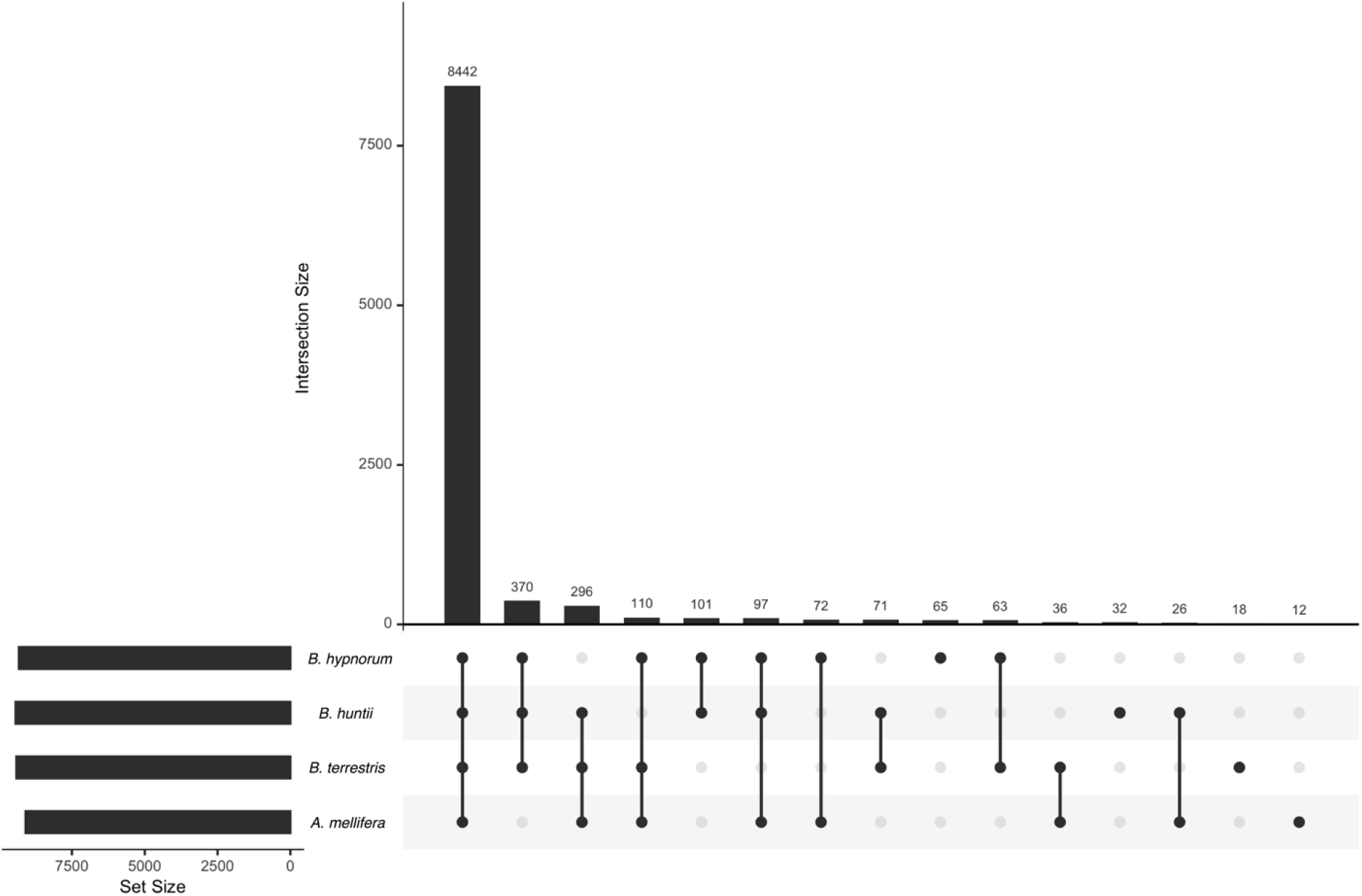
Orthogroup barplot of annotated genes for *B. huntii, B. hypnorum, B. terrestris, A. mellifera*. Size of bar represents the number of orthogroups identified for each taxon, shared between taxa, or unique to a taxon (i.e., species-specific orthogroup).

As the market for mass produced bumble bees grows in tandem with market needs, their domestication can be improved by genetic data and genomic resources. This is especially true for regulatory agencies and policy makers who make decisions on the importation of bumble bees across state, provincial, and national borders. Many of these decisions are driven in part by concerns on the movement of pathogens across bee species, hybridization, and increased interactions amongst native and invasive species. Even within species, there is concern about the use and escape of regional genotypes that could contribute to outbreeding depression among native locally adapted *Bombus* populations (Lozier *et al*. 2015). Furthermore, while *B. huntii* populations in the USA appear to be abundant and genetically diverse (Koch *et al*. 2015, 2018), populations in Mesoamerica are projected to be vulnerable to climate variation that is expected to occur over the next 30 years (Martínez-López *et al*. 2021). Identifying genetic loci under selection in the context of environmental variability for *B. huntii* will be useful in managing the commercial populations for pollination in diverse agricultural systems (Jackson *et al*. 2020).

## Data availability

The *B. huntii* genome assembly has NCBI Bioproject ID #PRJNA858880 (ToLID: iyBomHunt1) in the Beenome100 umbrella BioProject #PRJNA923301. GenBank accession is GCA_024542735.1 and RefSeq accession is GCF_024542735.1. PacBio HiFi reads are available on the NCBI Sequence Read Archive (SRA) at accession SRX16249823. Illumina sequences used for HiC scaffolding are available on SRA at SRX16249961.

## Acknowledgments

The genome assembly was generated as part of the USDA-ARS Beenome100 Initiative (https://www.beenome100.org/). This research used resources provided by the SCINet project of the USDA-ARS project number 0500-00093-001-00-D and the Tropical Pest Genetics and Molecular Biology project number JDL and JBUK were supported by the National Science Foundation DEB-2126418 and 2126417. The authors thank the members of the USDA-ARS Beenome100 and Ag100Pest Team for sequencing and analysis support. We also thank Tien Lindsay for her support in bombiculture and specimen curation. All opinions expressed in this paper are the authors’ and do not necessarily reflect the policies and views of USDA. Mention of trade names or commercial products in this publication is solely for the purpose of providing specific information and does not imply recommendation or endorsement by the USDA. The USDA is an equal opportunity provider and employer.

